# Genome assembly and annotations of a Cyprinidae fish, *Sinibrama wui*

**DOI:** 10.1101/2025.09.11.675668

**Authors:** Yanxuan Zhang, Jiaxiang Liu, Jinkai Liu, Dechun Zhang, Le Wang, Zhenzhen Xie

## Abstract

*Sinibrama wui* is a freshwater fish belonging to the Cyprinidae family, primarily found in the middle and upper Yangtze River of China. However, no reference genome sequence for *S. wui* has yet to be established. The present study presents a high-quality, chromosome-level genome assembly of *S. wui* through PacBio Biosciences sequencing and Hi-C technologies. The genome size was 1028.95 Mb, with a contig N50 length of 37.21 Mb and scaffold N50 of 40.74 Mb. We assembled 124 contigs, within which 68 were organized into 24 chromosomes. In total, 25,967 protein-coding genes were predicted, with 99.10% functionally annotated. We further analyzed the evolutionary relationships, gene families expansion, and positive selection of *S. wui*. Our data showed that *S. wui* diverged from *Megalobrama amblycephala* approximately 10.1 million years ago, and the expanded gene families had enriched in the lipid metabolism and nervous system. These datasets offer valuable genomic resources for future investigations studies on the genetic conservation, selective breeding, and evolution of *S. wui*.

## Introduction

The genus *Sinibrama* belongs to the cyprinid subfamily Cultrinae and represents a unique and economically important group of freshwater fish in China. *Sinibrama* species are typical potamodromous fishes widely distributed in the middle and upper reaches of the Yangtze River. It comprises four species: *S. macrops, S. wui, S. melrosei*, and *S. taeniatus*, respectively (Chen 1998). These species, characterized by their small size, fast growth, and rapid reproduction, hold immense value both economically and ecologically (Luo et al. 2014). However, in recent years, the wild populations of *Sinibrama* species have suffered significant declines, largely due to dam construction, the introduction of invasive or exotic species, overfishing, and environmental pollution (Zhao et al. 2018).

In response to these threats, efforts to artificial breeding and cultivation of these species have become crucial (Shi et al. 2023). A fundamental component of this conservation strategy is the accessibility of genomic resources. Unfortunately, there have been limited genomic resources reported for these species. To date, only complete mitochondrial genome sequences of *S. macrops* (Ai et al. 2013) and *S. taenictus* (Li et al. 2016) have been published. The scarcity of genomic resources has constrained genetic studies in these species, emphasizing the critical need for the development of these resources.

In this study, we present the first high-quality, chromosome-level genome assembly of *S. wui*, achieved through PacBio HiFi and Hi-C sequencing technologies. Our reference genome not only fills a critical gap in genomic resources but also serves as a robust foundation for investigations into genetic conservation, selective breeding, and evolution of *S. wui*.

## Materials and Methods

### Sample collection and DNA extraction

A wild male *S. wui* individual captured from the Chongyang River (26.6418°N, 118.1777°E) in Nanping, Fujian Province, China, was used for genome sequencing and assembly. The muscle tissue was used for genomic DNA extraction using a CTAB DNA extraction method (Chakraborty et al., 2020). The quality and quantity of the extracted DNA were examined using a NanoDrop 2000 spectrophotometer (NanoDrop Technologies, Wilmington, DE, USA) and a Qubit 3.0 Fluorometer (Invitrogen, USA). Nine distinct tissues (muscle, liver, skin, gill, gonad, intestines, kidney, eye, and brain) were collected for RNA extraction. All samples were immediately frozen in liquid nitrogen and then stored at −80 °C.

### Library construction and sequencing

We performed three sequencing techniques: Illumina, PacBio HiFi, and Hi-C, for *de novo* genome assembly of *S*.*wui*.

The DNA extracted from the muscle was first used for construction of a 300-bp insert library, which was further sequenced using Illumina Novaseq 6000 platform. In addition, 10 micrograms (ug) of genomic DNA were used for the construction of a 20-kb long fragment insert library using the SMRTbell Express Template Prep Kit 2.0 (Pacific Biosciences, USA). The library was further sequenced in circular consensus sequence (CCS) on the Pacific Biosciences Revio platform. The raw data was converted into high-precision HiFi reads using the CCS software (v.6.3.0, https://github.com/pacifcbiosciences/unanimity) (parameters: -min -Passes 3).

Then, muscle tissue of *S. wui* was used for Hi-C library construction, according to a previous study (Belaghzal et al. 2017). In brief, a white muscle sample of *S. wui* was dissected and cross-linked using 4% formaldehyde. The fixed sample was then lysed to isolate the nuclei. Subsequently, chromatin was digested with restriction enzyme MboI (NEB, USA). The digested DNA was then treated with end repairing, biotin labeling, and ligation of blunt-end fragments with T4 DNA ligase. After ligation, cross-linked products were reversed using 200 µg/mL proteinase K (Thermo, Shanghai, China) at 65 °C overnight. DNA was then purified using QIAamp DNA Mini Kit (Qiagen) according to the manufacturer’s instructions. The purified DNA was further sheared to a length of 300–500 bp and used for DNA sequencing library. The Hi-C library was quantified and sequenced on the MGI-SEQ2000 platform in 150PE mode.

For gene annotation, transcriptome sequencing was performed with nine tissues (liver, gill, intestines, kidney, brain, eye, gonad, skin, and muscle) of *S. wui*. Total RNA was extracted with the TRIzol reagent (Invitrogen, Waltham, MA, USA) according to the manufacturer’s instructions. The concentration and integrity of total RNA were estimated using an Agilent 2100 Bioanalyzer (Agilent Technologies, Santa Clara, CA, USA) and ethidium bromide staining of 28S and 18S ribosomal bands on a 1% agarose gel, respectively. The verified RNA samples were equally pooled for the RNA library construction and sequencing. In brief, the full-length cDNA was synthesized using the SMARTerTM PCR cDNA Synthesis Kit (Takara Biotechnology, Dalian, China). The SMRTbell library was constructed with the Pacific Biosciences DNA Template Prep Kit 2.0. The library was assessed using a Qubit 3.0 Fluorometer (Life Technologies, Carlsbad, CA, USA) for quantification and a 2100 Bioanalyzer system (Agilent Technologies, CA, USA) for quality analysis. Subsequently, SMRT sequencing was conducted with the PacBio Sequel II platform by Frasergen Bioinformatics Co., Ltd. (Wuhan, China).

### Genome survey and assembly

The size of the *S. wui* genome was estimated through K-mer analysis conducted with GCE (Liu et al. 2013) (https://github.com/fanagislab/GCE).

Initially, the raw Illumina sequencing reads underwent quality filtering using SOAPnuke (v2.1.0) (Chen et al. 2018) (main parameters: -lowQual=20, -nRate=0.005, -qualRate=0.5, other parameters as default). Subsequently, the quality-filtered reads were utilized to compute the K-mer frequency with a k-mer size 17 using Jellyfish (v. 2.2.10) (Marçais and Kingsford 2011). Then, the Pacbio HiFi reads were used to assemble contigs using the program Hifiasm v0.19.5 (Cheng et al. 2022) with 13 parameter. The reads obtained from the Hi-C library sequencing were aligned to the genome using Juicer (https://github.com/aidenlab/juicer, v3) with default parameters. Paired reads aligned to different contigs were employed to construct the Hi-C association scaffold. Furthermore, we utilized the 3D-DNA pipeline to cluster and orient the contigs to build a genome-wide chromosome interaction matrix (Kajitani et al. 2014). Finally, JuiceBox (v1.6.2) (Durand et al. 2016) was used to manually correct the assembly errors.

### Annotation of repetitive sequences

Two methods were combined to predict repetitive sequences in the genome: homology-based and *de novo* prediction approaches. For the homology-based analysis, known transposable elements (TEs) within the *S. wui* genome were identified using RepeatMasker (v4.1.2) (http://www.repeatmasker.org) with the Repbase library (v21.12) (Jurka et al. 2005). Repeat Protein Mask (v1.36) (http://www.repeatmasker.org) searches were also performed using the TE protein database as a query library. For *de novo* prediction, we constructed a *de novo* repeat library of the *S. wui* genome using RepeatModeler (v2.0.2a) and LTR_FINDER (v1.0.5) (Xu and Wang 2007). RepeatMasker was then used to search and classify the repeat regions against this newly constructed repeat library. Tandem Repeat Finder (TRF) (Benson 1999) was utilized to identify tandem repeats, while RepeatMasker was employed to identify non-dispersed repeat sequences. Finally, we merged the libraries constructed separately based on the two methods and subsequently utilized RepeatMasker to identify the repetitive sequences as a unified dataset.

### Gene Prediction and Functional Annotation

To predict protein-coding genes within the assembled genome of *S. wui*, we employed three distinct strategies: homology-based, *de novo*-based, and transcriptome sequencing-based protocols. Initially, protein sequences retrieved from Ensembl (Flicek et al. 2014) for *Megalobrama amblycephala, Onychostoma macrolepis, Danio rerio*, and *Sinocyclocheilus grahami* were aligned to the genome sequences of *S. wui* for homology annotation. Subsequently, the program Exonerate (v2.2.0) (Slater and Birney 2005) was used to execute homology-based gene prediction. Next, we employed Augustus (v3.3.1) (Stanke et al. 2006) and Genescan (v1.0) (Burge and Karlin 1997) for *de novo* gene prediction. Additionally, the identification of protein-coding genes based on assembled transcripts was carried out using GMAP (Wu and Watanabe 2005) as the third strategy. Following the initial predictions, PASA (Haas et al. 2003) was used to further refine the gene structures. Finally, MAKER (v3.00) (Holt and Yandell 2011) was employed to integrate the prediction outcomes from the three methods, resulting in a finalized set of non-redundant gene models.

Each of the predicted protein-coding genes underwent a rigorous functional annotation process, mapping to a variety of public databases. These included the non-redundant protein database (NR) (http://ftp.ncbi.nih.gov/blast/db/FASTA/nr.gz), TrEMBL (Holt and Yandell 2011), InterPro (Mitchell et al. 2015), SwissProt (Bairoch and Apweiler 2000), GO Ontology (GO) (Ashburner et al. 2000), and the Kyoto Encyclopedia of Genes and Genomes (KEGG) (Kanehisa and Goto 2000) databases. The annotation process was performed using BLASTP, selecting the best matching hit for each gene with an e-value threshold of 1e-5. The protein domains of predicted protein coding genes were annotated by employing PfamScan (Mistry et al. 2007) and InterProScan (v5.35–74.0) (Jones et al. 2014), based on InterPro (Blum et al. 2021) protein databases. Additionally, motifs and domains within individual gene models were identified through mapping to PFAM (Finn et al. 2008) databases.

### Gene family evolution and phylogenetic relationships

To discern the gene families for reconstructing phylogenetic relationship, we compared the predicted protein coding genes of *S. wui* with those of 15 other teleosts, including *D. rerio* (NCBI: GCA_000002035.4) (Howe et al. 2013), *M. amblycephala* (NCBI: GCA_018812025.1) (Liu et al. 2021), *S. grahami* (NCBI: GCA_001515645.1) (Yang et al. 2016), *Cyprinus carpio* (NCBI: GCA_004011595.1) (Xu et al. 2019), *Carassius auratus* (NCBI: GCA_003368295.1) (Chen et al. 2019), *Sinocyclocheilus anshuiensis* (NCBI: GCA_001515605.1) (Yang et al. 2016), *Sinocyclocheilus rhinocerous* (NCBI: GCA_001515625.1) (Yang et al. 2016), *Ctenopharyngodon Idella* (NCBI: GCA_019924925.1) (Wu et al. 2022), *Ancherythroculter nigrocauda* (NCBI: GCA_036281575.1) (Zhang et al. 2020), *Astyanax mexicanus* (NCBI: GCA_000372685.2) (Warren et al. 2021), *Ictalurus punctatus* (NCBI: GCA_001660625.1) (Liu et al. 2016), *Oreochromis niloticus* (NCBI: GCA_922820385.1) (Etherington et al. 2022), *Gasterosteus aculeatus* (NCBI: GCA_016920845.1) (Nath et al. 2021), *Oryzias latipes* (NCBI: GCA_000151825.1) (Kasahara et al. 2007), and *Lepisosteus oculatus* (NCBI: GCA_000242695.1) (Inoue et al. 2003). These sequences were reduced to create a nonredundant protein dataset by preserving the longest predicted isoform for each gene. Orthologous groups of protein-coding genes across the 16 species were identified using OrthoFinder2 (v2.5.4) (Emms and Kelly 2019) with default parameters.

To reconstruct the phylogenetic relationships between *S. wui* and the aforementioned 15 fish species, we identified and extracted 257 highly conserved single-copy orthologous genes across the 16 selected species. The protein sequences of the single-copy orthologous genes were aligned using the MUSCLE (v3.8.31) (Edgar 2004) program. Subsequently, the corresponding coding DNA sequence (CDS) alignments were generated and concatenated with the guidance of the protein alignment. The phylogenetic tree was constructed using RAxML (v8.2.12) (Stamatakis 2014) through the maximum likelihood method. *L. oculatus* was used as the outgroup in the analysis. The programs R8s (v1.81) (Sanderson 2003) and the MCMCTree, implemented in the PAML (v4.10.0) (Yang 2007) packages, were used to estimate the divergence time among species, with default parameters.

### Expansion and contraction of gene families

Based on the identified gene families, the constructed phylogenetic tree, and the predicted divergence time of the studied fish, we used CAFE (v4.2.1) (De Bie et al. 2006) to detect evidence of gene expansions and contractions within gene families.

### Positively selected genes

Based on the phylogenetic tree, we estimated the ratio (ω) of nonsynonymous (Ka) to synonymous (Ks) nucleotide substitutions using the PAML (v4.10.0) (Yang 1997) package. This analysis allows us to examine the selective constraints on the individual candidate genes. The single-copy orthologous genes across the studied species were initially aligned using the program Muscle (v3.8.31) (Edgar 2004) and subsequently refined using the program trimAl v1.4 (Capella-Gutiérrez et al. 2009). For each gene, we compared a series of evolutionary models within the likelihood framework for each gene using the species trees. A branch-site model was used to detect the average value of ω across the tree (ω0), ω of the appointed branch under investigation (ω2), and ω of all the remaining branches (ω1).

## Results and Discussion

### Genome Assembly and Assessment

The estimated genome size of *S. wui* was 980.44 Mb, featuring a heterozygosity rate of 0.66% and comprising repeat sequences accounting for 54.72% (refer to supplementary table S1 and fig. S1a, Supplementary Material online).

A total of 45.14 Gb (~43.9× depth) PacBio sequencing data with a mean length of approximately 18.5 kb and an N50 length of 18.7 kb were generated and used for genome assembly (refer to supplementary table S2, Supplementary Material online). This resulted in 138 contigs, with a combined length of 1.07 Gb and a contig N50 of 37.21 Mb (refer to Table 1 and fig. 1a). The genome size of *S. wui* was shorter than the genome of *Megalobrama amblycephala* (1.11Gb) (Liu et al. 2021), but longer than the genome of *Ancherythroculter nigrocauda* (1.04Gb) (Zhang et al. 2020). Additionally, chromosome genome assembly is crucial for genome comparison and evolutionary studies (Jaillon et al. 2004). In the present study, approximately 107.91 Gb (~106.0× depth) clean reads were used to construct the chromosome-level genome of *S. wui* (refer to supplementary table S2, Supplementary Material online). We observed that 719.9 million paired-end reads from the Hi-C libraries were aligned to the assembled contigs, comprising 99.67% of the total cleaned paired-end reads (refer to supplementary table S3, Supplementary Material online). According to the Hi-C results, a total of 68 contigs were anchored into 24 pseudochromosomes (Chr) with a total length of 1.02 Gb, a contig N50 length of 37.21 Mb, and a scaffold N50 length of 40.74 Mb (refer to supplementary table S4 and fig. S1b, Supplementary Material online).

**Table 1.** Statistic of *Sinibrama Wui* Genome Assembly and Annotation Data.

| Genome Assembly |  |  |
| --- | --- | --- |
| Assembly | Genome size (bp) | 1,065,979,054 |
|  | Number of scaffolds | 94 |
|  | Number of contigs | 138 |
|  | Scaffold N50 (bp) | 39,564,191 |
|  | Contig N50 (bp) | 37,212,952 |
|  | GC (%) | 37.3 |
| BUSCO (% of total BUSCOs) | Complete | 942 (98.8%) |
|  | Single-copy | 923 (96.8%) |
|  | Duplicated | 19 (2.0%) |
|  | Fragmented | 5 (0.5%) |
|  | Missing | 7 (0.7%) |
| Repetitive elements | DNA (bp) | 437,493,010 (42.52%) |
|  | LINE (bp) | 65,137,353 (6.33%) |
|  | SINE (bp) | 3,466,455 (0.34%) |
|  | LTR (bp) | 128,715,757 (12.51%) |
|  | Others | 12,338 (0.00%) |
|  | Unknown (bp) | 41,218,883 (4.01%) |
|  | Total (bp) | 607,375,105 (59.03%) |
| Gene annotations (% of all genes) | Swissprot | 22,277 (85.79%) |
|  | NR | 25,720 (99.05%) |
|  | KEGG_ALL | 25,182 (96.89%) |
|  | TrEMBL | 24,903 (95.90%) |
|  | GO | 20,160 (77.64%) |
|  | KEGG_KO | 16,684 (64.25%) |
|  | InterPro | 22,036 (84.86%) |
|  | Total | 25,967 (100%) |

**Fig. 1.**
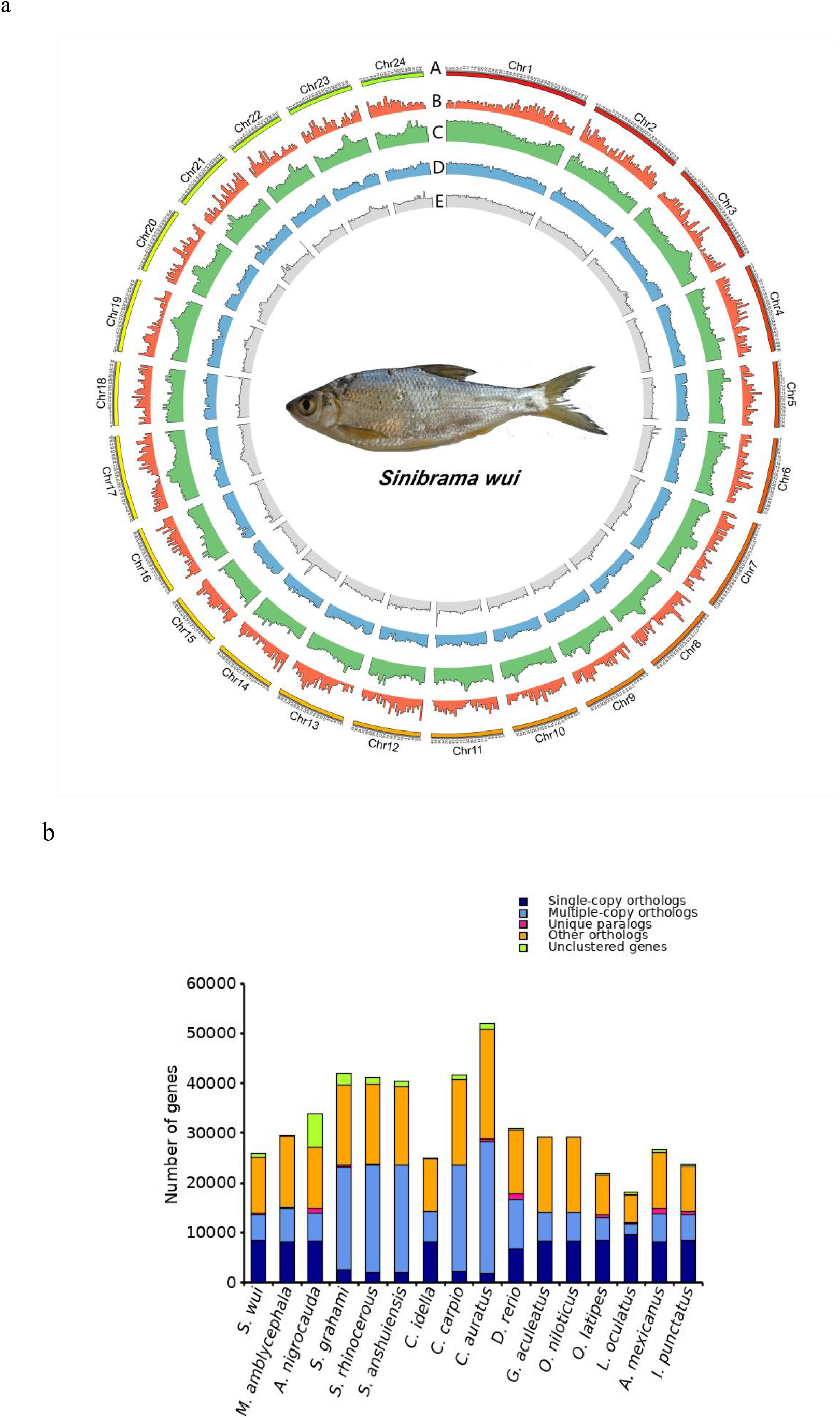
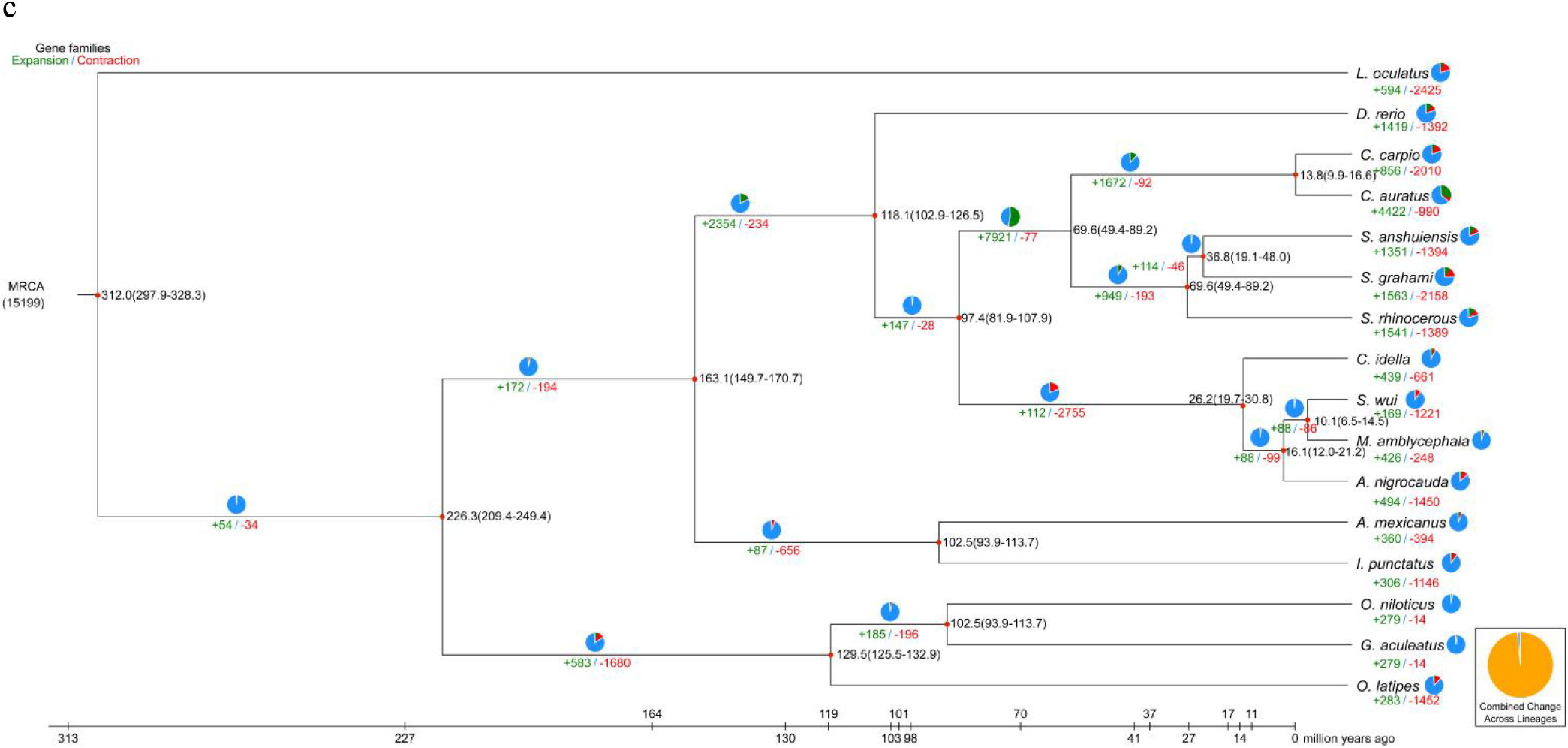
(a) Features of *Sinibrama wui* genome. (b) Comparison of the number of homologous genes. (c) *S*.*wui* diverged from other species and their phylogeny.

According to the Benchmarking Universal Single-copy Orthologs (BUSCO) results, a remarkable 98.8% (942) of the BUSCOs in the Actinoperygii_Odb10 database were identified as complete genes in the assembled genome. This includes 96.8% (923) complete and 2% (19) duplicated BUSCOs, as well as 0.5 % (5) fragmented BUSCOs (refer to Table 1).

### Gene Structure Prediction and Functional Annotation

Two methods were combined to predict repetitive sequences in the genome: homology-based and *de novo* prediction approaches. We observed that repetitive sequences were 634.55 Mb in length, accounting for 61.67% of the assembled genome. Meanwhile, transposable elements (TEs) represented 59.03% of the assembled genome, with a length of 607.38 Mb, which divided into four groups: DNA transposons (42.52%), long terminal repeats (LTRs, 12.51%), long interspersed elements (LINEs, 6.33%) and short interspersed nuclear elements (SINEs, 0.34%) (refer to supplementary table S5, Supplementary Material online).

In order to predict protein-coding genes within the assembled genome of *S. wui*, we employed three distinct strategies: homology-based, *de novo*-based, and transcriptome sequencing-based protocols. Based on *de novo* methods, a total of 26,286–28,719 genes were detected. A comparison with four selected teleost genomes of *Megalobrama amblycephala, Onychostoma macrolepis, Danio rerio*, and *Sinocyclocheilus grahami*, a total of 5,390–65,140 proteins were indicated. Based on transcriptome sequencing data, a total of 14,016 genes were detected. Finally, a total of 25,967 protein-coding genes, with an average gene length of 19,708.54 bp, were identified across the entire genome assembly. The number of predicted protein-coding genes in *S. wui* was high than those in *M. amblycephala* (23,696) (Liu et al. 2017) and *O. macrolepis* (24,770) (Sun et al. 2020), but lower than those in *A. nigrocauda* (34,414) (Zhang et al. 2020). Moreover, the average coding sequence (CDS) length, exon length, and intron length were 1,665.4, 246.74, and 2,052.36 bp, respectively (refer to supplementary table S6, Supplementary Material online and fig. 1a). The *S. wui* genome annotation included 3,219 (96%) complete copies, with 3,127 (94.6%) of BUSCO genes being complete and single copy, 47 (1.4%) being complete and duplicated, 51 (1.5%) being fragmented, and 84 (2.5%) being missed (refer to supplementary table S7, Supplementary Material online). Additionally, for the predicted non-coding genes, 2,496 microRNAs (miRNAs), 18,709 transfer RNAs (tRNAs), 10,565 ribosomal RNAs (rRNAs), and 941 small nuclear RNAs (snRNAs) were also identfied in the genome of *S. wui* (refer to supplementary table S8, Supplementary Material online).

Each of the predicted protein-coding genes underwent functional annotation through mapping to various public databases, including the non-redundant protein database (NR), TrEMBL, InterPro, SwissProt, GO Ontology (GO), and the Kyoto Encyclopedia of Genes and Genomes (KEGG) databases. As a result, out of the predicted protein-coding genes, approximately 25,732 (99.10%) were functionally annotated, with the Swissprot database annotated 22,277 (85.79%), the NR database annotated 25,270 (99.05%), the KEGG database annotated 25,182 (96.89%), the TrEMBL database annotated 24,903 (95.90%), the Go database annotated 20,160 (77.64%), the KOG database annotated 16,684 (64.25%), and the InterPro database annotated 22,036 (84.86%), respectively (refer to Table 1).

### Gene family evolution and phylogenetic relationships

A total of 25,166 (96.92%) out of the 25,967 protein-coding genes were clustered into 18,833 gene families in *S. wui* (refer to fig. 1b), including 257 single-copy gene families shared with other fish species and 66 unique gene families (refer to supplementary table S9, Supplementary Material online). On average, each orthologous group contained 1.33 genes. Additionally, 801 genes were unclustered, indicating that they may be species-specific. Furthermore, we compared the gene families among the four related species (*S. wui, M. amblycephala, O. macrolepis*, and *S. grahami*), and found that 15,958 orthologous gene families were shared among the four species, of which 257 gene families were specific to *S. wui* (refer to supplementary fig. S2, Supplementary Material online). The maximum likelihood phylogenetic tree suggests that *S. wui* and *M. amblycephala* diverged from their common ancestor approximately 6.5 to 14.5 million years ago (mya), after diverging from *A. nigrocauda* around 16.1 mya (refer to fig. 1c).

Understanding the expansion and contraction of gene families is essential for unraveling phenotypic diversity and environmental adaptation (Harris and Hofmann 2015; Xie et al. 2022). In comparison with the selected teleost species, a total of 169 and 1,221 gene families showed evidence of expansion and contraction, respectively, in *S. wui* (refer to fig. 1c). Among the gene families displaying evidence of expansion,19 KEGG pathways, and 20 GO terms were significantly enriched (refer to supplementary fig. S3, Supplementary Material online). Additionally, a total of 251 positively selected genes were identified within the *S. wui* genome. These enriched GO terms were found to be associated with functions related to “binding” and “cell part”. Furthermore, the KEGG enrichment analysis highlighted that the expanded gene families were significantly enriched in pathways associated with lipid metabolism - arachidonic acid metabolism (ko:00590) and nervous system - serotonergic synapse (ko:04726) (refer to supplementary table S10, Supplementary Material online). It has demonstrated that the lipid metabolism in fish may have significance in inducing hepatic oxidative stress and inflammation (Liu et al., 2024). Besides, the nervous system in fish may have significance in the evolutionary of fish (Weil et al., 2018).

## Conclusion

*Sinibrama wui* is an economically important species in the middle and upper reaches of the Yangtze River. However, there have been limited genomic resources reported for these species. In the present study, we have presented the first high-quality, chromosome-level genome assembly of *S. wui* using PacBio sequencing and Hi-C technologies. The genome size was 1028.95 Mb, with a contig N50 length of 37.21 Mb and scaffold N50 of 40.74 Mb. Besides, 25,967 protein-coding genes were predicted, with 99.10% functionally annotated. The newly generated reference genome will serve as a robust foundation for studies on the genetic conservation, molecular breeding, as well as evolutionary studies of *S. wui*.

## Data Availability

All data supporting the funding of this study has been deposited in the NCBI Sequence Read Archive under the BioProject number PRJNA1090244. All of the raw sequencing data, including Illumina, PacBio HiFi reads, Hi-C, and transcriptome data, are also deposited under the same BioProject number. The genome assembly is deposited into NCBI under the accession number GCA _040182825.1. The genome annotation files are available in the Genome Warehouse in National Genomics Data Center, Beijing Institute of Genomics, Chinese Academy of Sciences / China National Center for Bioinformation, under accession number GWHETLY00000000.1 that is publicly accessible at https://ngdc.cncb.ac.cn/gwh..

Supplemental material is available at G3 online.

## Acknowledgments

ZX and LW designed and coordinated the project. XL wrote the manuscript. JXL collected and process the samples. JKL completed the sequencing and data processing. DZ performed genome assembly and annotation. LW participated in language editing. ZX reviewed the manuscript. All authors have read and approved the final manuscript.

## Conflictes of interest

The authors declare no conflict of interest.

## Author contributions

ZZX and LW conceived of the project. JKL and DCZ collected the sample. YXZ and JXL performed the molecular work. YXZ, JXL and DCZ analyzed the data. YXZ wrote the manuscript. All authors reviewed the manuscript.

